# PlasmidGPT: a generative framework for plasmid design and annotation

**DOI:** 10.1101/2024.09.30.615762

**Authors:** Bin Shao

## Abstract

We introduce PlasmidGPT, a generative language model pretrained on 153k engineered plasmid sequences from Addgene. PlasmidGPT generates *de novo* sequences that share similar characteristics with engineered plasmids but show low sequence identity to the training data. We demonstrate its ability to generate plasmids in a controlled manner based on the input sequence or specific design constraint. Moreover, our model learns informative embeddings of both engineered and natural plasmids, allowing for efficient prediction of a wide range of sequence-related attributes.

## Introduction

Plasmids are essential tools for genetic manipulation. They are commonly used as vectors for introducing new genetic material into organisms, which enables the study of gene function and the production of recombinant proteins^1^.Beyond basic research, plasmids are crucial in biotechnological applications, allowing for the development of vaccines and the engineering of microbial strains for biomanufacturing.

Despite its significance, design of plasmid DNA is still a labor-intense effort which requires manual inspection, annotation and combination of functional units. Furthermore, we lack a powerful tool to study the grammar inherent in the growing collection of engineered plasmid sequences, hindering their reusability and limiting our ability to standardize and automate plasmid design. Recently, generative models like Generative Pre-trained Transformers (GPT)^2^ have shown to be highly successful in modeling human language. Due to the similarity of human language and biological sequences like protein and DNA, researchers have adapted these modeling frameworks to synthesize novel proteins^3^ and more recently, to genomic sequences^4,5,6^.

Here we introduce PlasmidGPT, a generative framework for designing and annotating plasmid DNA sequences (Fig. 1a). Our framework is built on a decoder-only transformer model that is pretrained on 153k plasmid sequences from Addgene, a public repository for engineered DNA sequences. PlasmidGPT generates novel plasmid sequences rather than replicating training data, and the generated plasmids have genetic part distributions similar to those of the training sequences. Conditional plasmid generation can be achieved by either providing a user-specified starting sequence or by fine-tuning the model for specific vector types. Finally, PlasmidGPT extends beyond generation to effectively predict features of both engineered and natural plasmids, highlighting its potential as a versatile tool for plasmid analysis. Our model is publicly available on GitHub: https://github.com/lingxusb/PlasmidGPT

**Figure 1.**
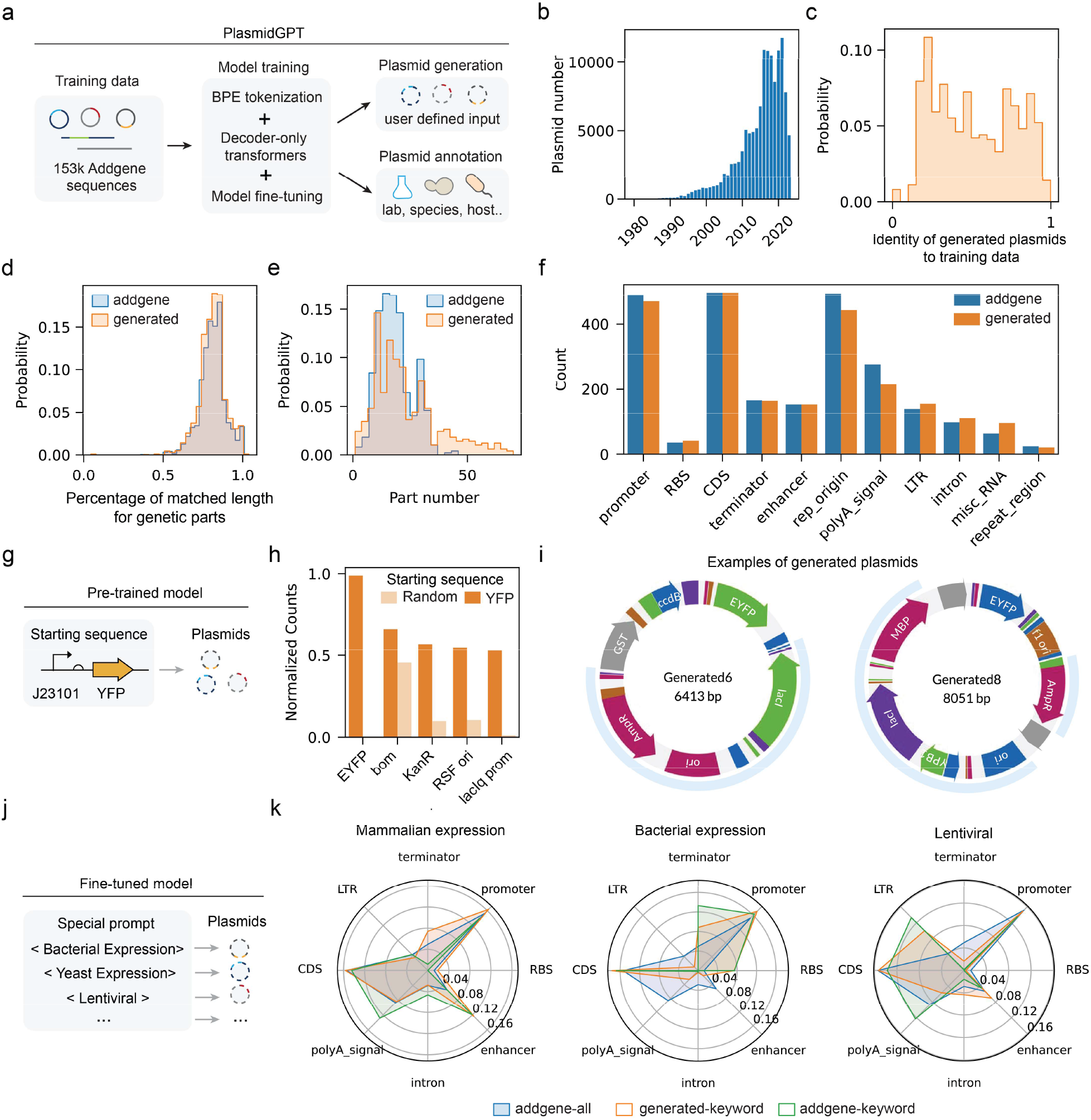
Generation of plasmid sequences using a language model. **a)** Overview of model training and application. **b)** Distribution of plasmid publication dates across the training dataset. **c)** Histograms showing the distribution of the identity of the generated plasmid sequences (n = 1,000) compared to the training dataset. The closet match to the training dataset was identified by BLAST analysis^8^. Histograms showing the distribution of the percentage of matched length for genetic parts **(d)** and the number of parts **(e)** for the training dataset (n = 153,208) and the generated plasmid sequences (n = 1,000). Genetic parts were identified using pLannotate^9^. **f)** Counts for the most frequent parts for randomly sampled training and generated sequences (n = 1,000). **g)** Conditional generation of plasmid sequence from a YFP expression cassette. **h)** Counts for the most frequent parts in the conditional generated sequences, compared with sequences generated with random 4-nt starting sequences (n = 1,000). **i)** Two generated sequences starting from the YFP expression cassette. Parts longer than 300 nt are annotated, and incomplete parts are represented in gray. Shaded areas indicate the overlap with the closest match in the training dataset. **j)** Finetuned PlasmidGPT model enables generation of plasmid sequences based on specific prompts. **k)** Part frequences are shown for randomly sampled training dataset (blue, n = 2,000), training samples related to specific keywords (green, n = 200) and generated sequences with specific keywords (orange, n = 100).

## Results

We collected the plasmids sequences longer than 2k base pair (bp) from Addgene to train our generative model (Fig. 1b). The Byte Pair Encoding (BPE) algorithm was applied for tokenization^7^, which breaks down the sequences into subunits. The model we used is a decoder-only transformer model consisting of 12 layers and 110 million parameters. After model training, we generated 1k plasmid sequences using random starting sequences and a temperature of 1.0, ensuring a high level of diversity among them. We found that the average length of generated sequences is slightly longer than the training sequences (Supplementary Fig. 1). To compare the similarity between the two, we used BLAST analysis^8^ to search the generated sequences against the training data and calculated the relative size of the overlap region (identity). Our results show that a large proportion of the generated plasmids have an identity lower than 0.5, indicating that they are substantially different from the training sequences (Fig. 1c).

Next, we used pLannote^9^ to annotate the genetic parts in both the generated and training sequences. The pLannote software includes a database of commonly used components, including promoters, ribosome binding sites (RBS), gene coding regions, and replication origins. For each plasmid, pLannote provides the annotated parts along with their matched percentage to the feature library. The generated and training sequences show similar distributions for the matched percentage of annotated parts (Fig. 1d). The part numbers for the generated sequences exhibit a broader distribution than those in the training data (Fig. 1e), though the median values of the two distributions are close with each other (17 vs. 19). We then compared the frequencies of different genetic parts between the generated and training sequences (Fig. 1f). The relative abundances of these genetic parts were also similar across both datasets, with promoters, gene coding regions (CDS), and replication origins showing the highest frequencies. These findings suggest that our model produces sequences that share similar structural characteristics with known plasmid sequences.

An important feature of generative models is their ability to generate sequences conditionally based on user’s input, just as how GPT models can write an article from a starting sentence. To explore this ability of the plasmidGPT model, we generated plasmid sequences using a constitutively expressed YFP cascade^10^ as the input and annotated the resulting sequences (Fig. 1g, Supplementary Table 1). We found that these sequences were more likely to contain components like the bom site, KanR (kanamycin resistance gene), and RSFi (replication origin), compared to sequences generated with random primers (Fig. 1h). We also presented two examples of generated plasmids (Fig. 1i). While these sequences share some overlap with the training data, they also include novel features, such as the *ccdb* gene which is widely used as part of a positive selection system for molecular cloning^11^.

To improve control over the plasmid generation process, we fine-tuned the PlasmidGPT model using vector type information as input prompt (Fig. 1j). We selected the 10 most common vector types in the training data and encoded each type as a special token (Supplementary Fig. 2). After model fine-tuning, we generated 1,000 sequences for each prompt. Our results revealed that generate sequences recapitulated the distinctive distribution of genetic parts associated with each vector type in the training data (Fig. 1k and Supplementary Fig. 3). For example, sequences generated with the “mammalian expression” prompt showed a high proportion of enhancers, which are crucial for gene expression in mammalian cells. In contrast, sequences generated with the “bacterial gene expression” prompt had high proportion of promoters but few polyA signals, consistent with the requirements for bacterial gene expression.

Sequence embeddings are numerical representations of input data produced by language models. They have been widely used to predict functional properties and understand the underlying structure of input sequences. We explored whether the embeddings generated by PlasmidGPT could provide meaningful insights into the functionality of plasmid sequences. For example, associating engineered plasmid sequences with lab of origins could help promote responsible biotechnology innovation^12–14^. Here we utilized PlasmidGPT’s learned embeddings to predict four attributes related to engineered plasmid sequences, including lab of origin, vector type, species and growth strains (methods). For each attribute, we trained a separate single-layer neural network using the sequence embeddings as input and evaluated its performance using 5-fold cross-validation tests (Fig. 2a).

**Figure 2.**
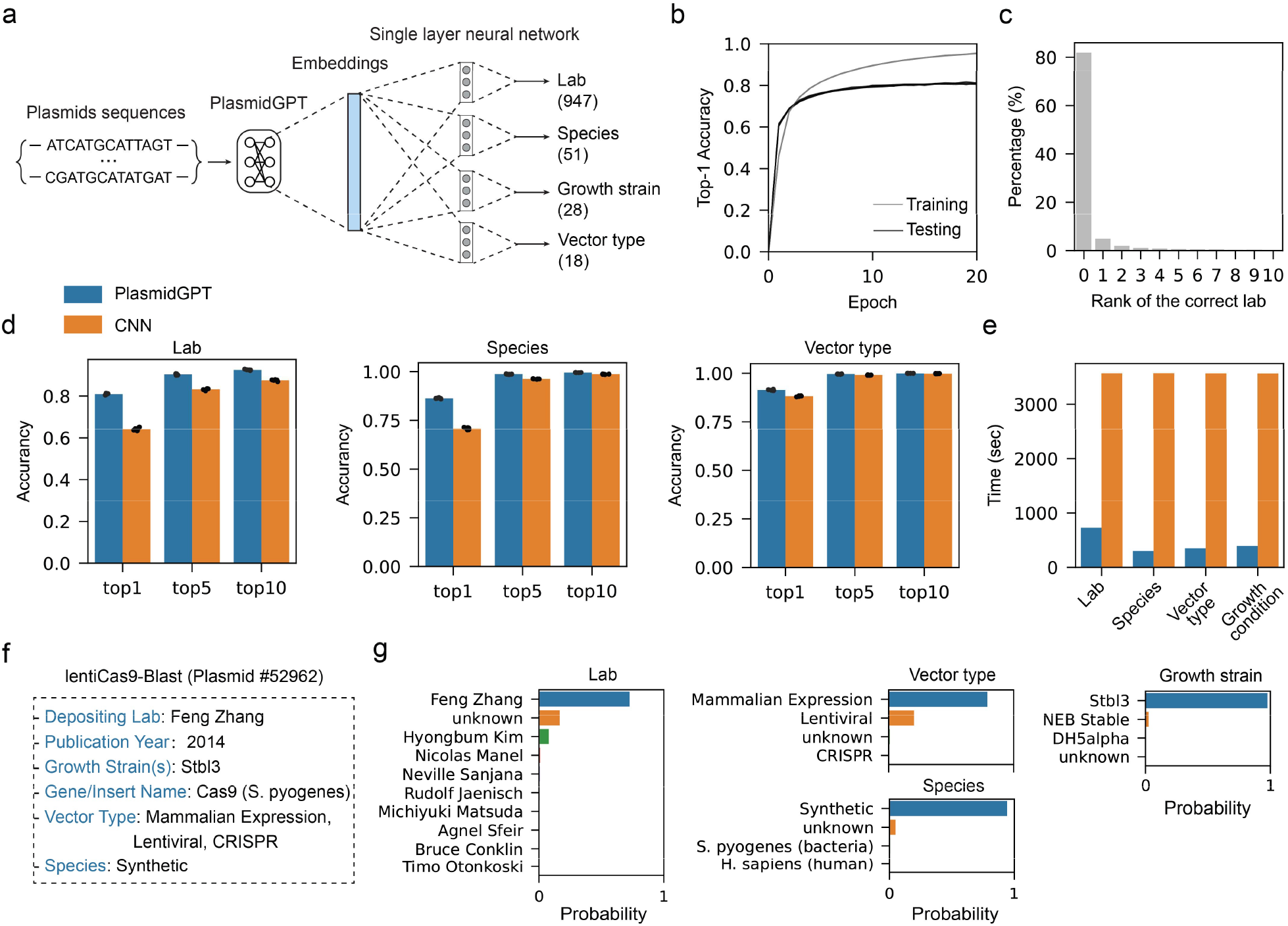
Annotation of engineered plasmids. **a)** Prediction of attributes based on PlasmidGPT model embeddings. Numbers present counts of unique labels. **b)** Top 1 accuracy for the lab of origin prediction task on the training and testing datasets as a function of training epochs. Results from 5-fold cross validation tests are shown (n = 5). **c)** Distribution of the rank of the correct lab among the model predictions. **d)** Performance of PlasmidGPT and CNN model on the predictions of lab of origin, species and vector types. We used 5-fold cross validation tests to evaluate model performance (n = 5), and error bars denote standard derivatives. **e)** Computational time required by PlasmidGPT and the CNN model for the prediction of different attributes. **f)** An example plasmid (#52962) in the test dataset was used to illustrate model performance. **g)** Top predictions for the example plasmid.

For the lab of origin prediction task, both validation and training loss plateaued after epoch 20 (Fig. 2b). The model achieved a top 1 accuracy of 81% and a top 10 accuracy of 92%, both higher than the best values reported in previous works (76% top 1 accuracy and 90% top 10 accuracy)^13,14^ (Fig. 2c). Additionally, we trained a convolutional neural network (CNN) based on the one-hot encoded nucleotide sequences, following the work by Nielsen et al^12^ but with a larger context window (16k bp). We found that our single-layer neural network model outperformed the CNN model by 17% percent for the top 1 accuracy (Fig. 2d), demonstrating its ability to capture the intrinsic structure of plasmid sequences. It also outperformed the CNN model on the species and vector type prediction tasks, with the improvement of 16% and 3% respectively. PlasmidGPT and CNN model achieved similar performance for vector type prediction (Supplementary Fig. 4). Using plasmid #52962^15^ from the test dataset as an example, our trained model correctly predicted all related variables (Fig. 2f and 2g). For the vector type, the right labels are within the top 4 predictions. Interestingly, *S. pyogenes* was among the top species predictions even the labeled species for this plasmid is “synthetic”. Furthermore, our model’s computational time is one order of magnitude lower than the vanilla CNN model (Fig. 2e), potentially making it more suitable for real-time applications.

To better understand how the trained model makes its predictions, we investigated the contribution of individual tokens for identifying the lab of origin. For this analysis, we focused on plasmids originating from the Feng Zhang lab. For each plasmid, one token was removed at a time from the tokenized sequence, and we then compared the activity of the output neuron corresponding to the Feng Zhang lab with that of the unaltered sequence. We found that most frequently occurring tokens have a mean deletion effect close to zero (Supplementary Fig. 5). However, a few tokens with high frequency showed a significant negative deletion effect, meaning that removing them severely affects the model’s ability to predict the Feng Zhang lab. One such token is 11824, harboring the puroR gene and EF-1*α* promoter, both of which are important for gene selection and expression. These findings highlight our model’s ability to identify key signatures in engineered sequences that are linked to specific labs.

We further evaluated our model’s performance on out-of-distribution (OOD) samples, which is crucial for understanding its capacity to handle unseen data. Our training and testing datasets have a cutoff date of February 2023. To conduct the OOD evaluation, we obtained a plasmid published in July 2023^16^ (Addgene #212888, Supplementary Fig. 6). Notably, our model accurately predicted key variables related to this plasmid, including lab of origin (Chris Voigt lab) and the growth strain (DH5alpha). The only variable that the model failed to predict is the species (*Salinispora Arenicola*), which was not included in the training labels. In this case, the model returns an “unknown” label and avoids making potentially misleading predictions.

Finally, we extended our modeling framework to the analysis of natural plasmids. Using sequences from the IMG/PR database^17^, we calculated sequence embeddings with the PlasmidGPT model. These embeddings were then used as input for a single-layer neural network to predict the host taxonomy of the natural plasmids. We compared this approach to a CNN model trained on one-hot encoded nucleotide sequences and both models were evaluated with 5-fold cross-validation tests. Despite being pretrained on engineered plasmid sequences, our PlasmidGPT model achieved comparable performance to the CNN model and required much less computational time (Supplementary Fig. 7), suggesting that it learns structural features that are shared between both natural and engineered plasmid sequences.

## Discussion

In this study, we leverage a generative language model to facilitate the design and analysis of plasmid sequences. By treating DNA sequences as a form of language, our approach enables more intuitive and efficient interaction with genetic information compared to the traditional labor-intensive process. However, our current model is not without limitations. First, it is important to note that the plasmids generated by PlasmidGPT are not yet comparable in complexity, functionality, or reliability to those produced by well-established methods like Cello^10,18^. Second, our model relies on tokenization of the input sequence, and the limited vocabulary may not fully capture the diversity of genetic parts. Despite these limitations, we believe that further developments of tools like PlasmidGPT have the potential to lower the technical barrier for sophisticated DNA design and provide novel insights into plasmid biology and evolution.

## Methods

### Model training and inference

Plasmid DNA sequences and their corresponding meta data were downloaded from Addgene (https://www.addgene.org/) with a time cutoff of February 2023. In cases there are multiple sequences related to one plasmid, the longest one was chosen for further analysis. Sequences shorter than 2kb were removed, resulting in a training dataset of 923M base pairs (bp). The Byte Pair Encoding (BPE) tokenizer^7^ was trained on all the plasmid sequences with a vocabulary size of 29,999. Given the circular nature of plasmids, we implemented data augmentation to expand our training dataset. For each tokenized plasmid sequence, we selected 10 random positions, and new sequences were constructed by placing the latter sequence after the position before the sequences preceding these positions. This procedure results in a ten-fold expansion of the training dataset.

We trained a decoder-only transformer model on the augmented dataset. The model consists of 12 layers with a dimension of 512, with 8 attention heads and a total of 110M parameters. The model training was conducted using the Adam optimizer, a learning rate of 0.0002 and a batch size of 1. For model inference, a temperature of 1.0 and a top k value of 50 were utilized to generate diverse sequences. A random token from the training dataset was selected as the starting sequence and the maximum context size was set to 200 tokens. For model training and inference, we used Nvidia’s A100 GPU (40GB) and 3090 Ti GPU (24GB) and the software packages PyTorch (version 2.1.1) and transformer (version 4.28.1).

### Model fine-tuning

To fine-tune the pretrained PlasmidGPT model, we used the top 10 vector types as key words. Since the number of plasmids for mammalian expression and bacteria expression is much higher than the other vector types, we subsampled these two categories to approximately 10,000 plasmids to balance the dataset. After tokenizing the plasmid sequences, we incorporated a special token at the beginning of each tokenized sequence to encode the corresponding vector type information. We fine-tuned the model using a learning rate of 0.0002 and a batch size of 1. The model was trained for a total of 10 epoch. When generating plasmid sequences specific to vector types, we used a temperature of 1.0 and top k value of 50. The repetition penalty was set to 1.0 and the maximum length was limited to 200 tokens. These settings were designed to balance creativity and coherence in the generated sequences.

### Prediction of attributes of plasmid sequences

We gathered metadata from the Addgene website for the plasmid sequences, including information about the lab of origin, vector type, species and growth strains. For the lab of origin, we only included the top 1000 labs and removed those with fewer than 10 sequences. This filtering process results in 947 unique lab labels, with the remaining plasmids labeled as “unknown”. In the case of species, we selected the top 100 species from the metadata and manually curated the labels by merging different names for the same species, which leads to 51 different species labels. For growth strains, we chose the top 30 strains and removed those with fewer than 10 plasmid sequences, resulting in 28 growth strain labels. For vector types, we selected the top 30 vector types and manually combined those with different names. This process gives us a total of 18 different vector types.

For the natural plasmids, we downloaded sequences from IMG/PR website (https://genome.jgi.doe.gov/portal/IMG_PR/IMG_PR.home.html). This dataset contains 279k natural plasmids with host taxonomy annotation. We selected all major bacterial phyla that had more than 10 associated plasmids, leading to 26 distinct host phyla.

To predict plasmid-related attributes, we used the tokenized plasmid sequences as model input to calculate their sequence embeddings (dimension 768). Then a single-layer neural network was trained (dimension: 128), with an input dimension of 768 and an output dimension corresponding to the number of classes for each attribute. The model was trained with a learning rate of 0.001 and a batch size of 64 for 20 epochs. 5-fold cross validation tests were used to evaluate the model performance: in each round of test, the data was randomly split into 5 folds, where 4 folds were used for training and the remaining fold for evaluation.

### CNN model for the prediction of plasmid attributes

We followed the Neilsen et al.^12^ to model plasmid sequences using CNN model. To further boost model performance, we used a context length of 16k bp, which is longer than the 8k bp in the original work. In brief, we did one-hot encoding of the raw DNA sequences with 0, 1, 2, 3, 4 encoding A, T, C, G and N respectively. The first layer is a 1D convolutional layer with 128 filters, each with a kernel size of 12. The convolution operation is followed by a max-pooling layer and batch normalization. The pooled output is flattened and passed through a fully connected layer with 64 nodes, followed by batch normalization. The final fully connected layer outputs predictions for different classes. The model’s parameters were optimized using the Adam optimizer with a learning rate of 0.001, and the loss was calculated using the cross-entropy function (*torch*.*nn*.*CrossEntropyLoss*). We employed 5-fold cross-validation to evaluate the model performance, and we calculated Top-1, Top-5, and Top-10 accuracies on the test set (without the “unknown” label).

## Code availability

Our trained model and inference codes are available from GitHub: https://github.com/lingxusb/PlasmidGPT

## Supplementary Figures

**Supplementary Figure 1.**
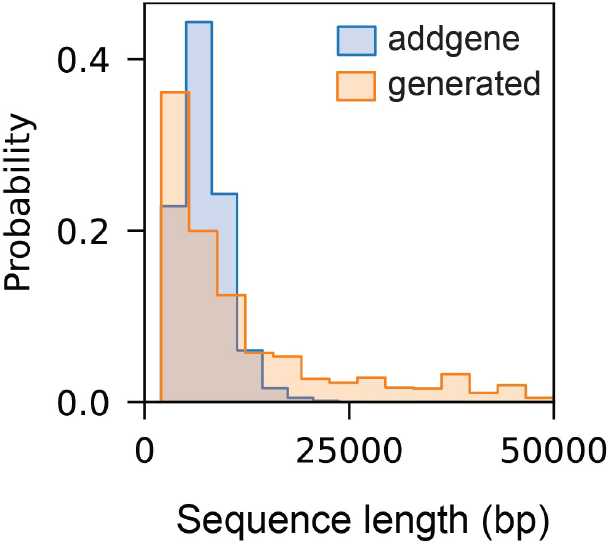
Sequence length distributions for the generated and training sequences. Sequence length of the generated sequences (n = 1000) versus the training dataset (n = 153,208).

**Supplementary Figure 2.**
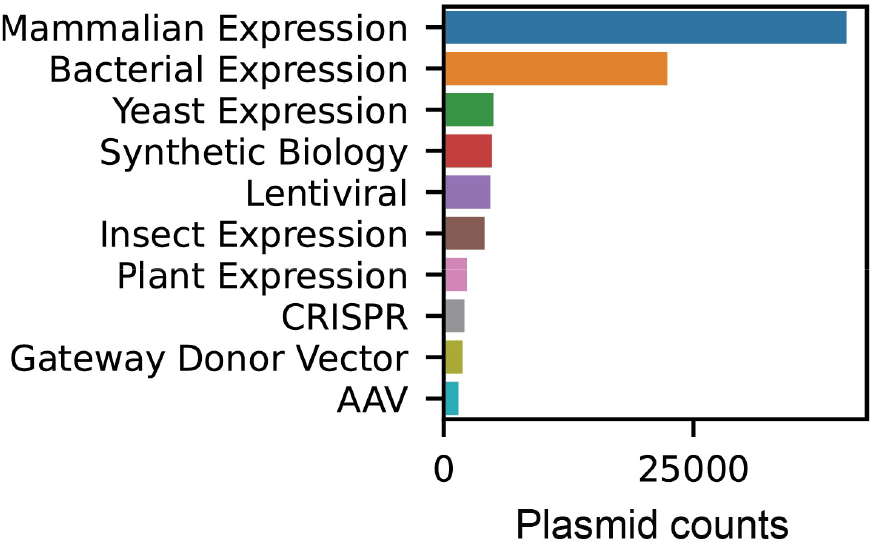
Top 10 vector types for the Addgene plasmids. Plasmid descriptions were retrieved from the addgene website(https://www.addgene.org/). Plasmid counts for the 10 most frequent vector types are reported.

**Supplementary Figure 3.**
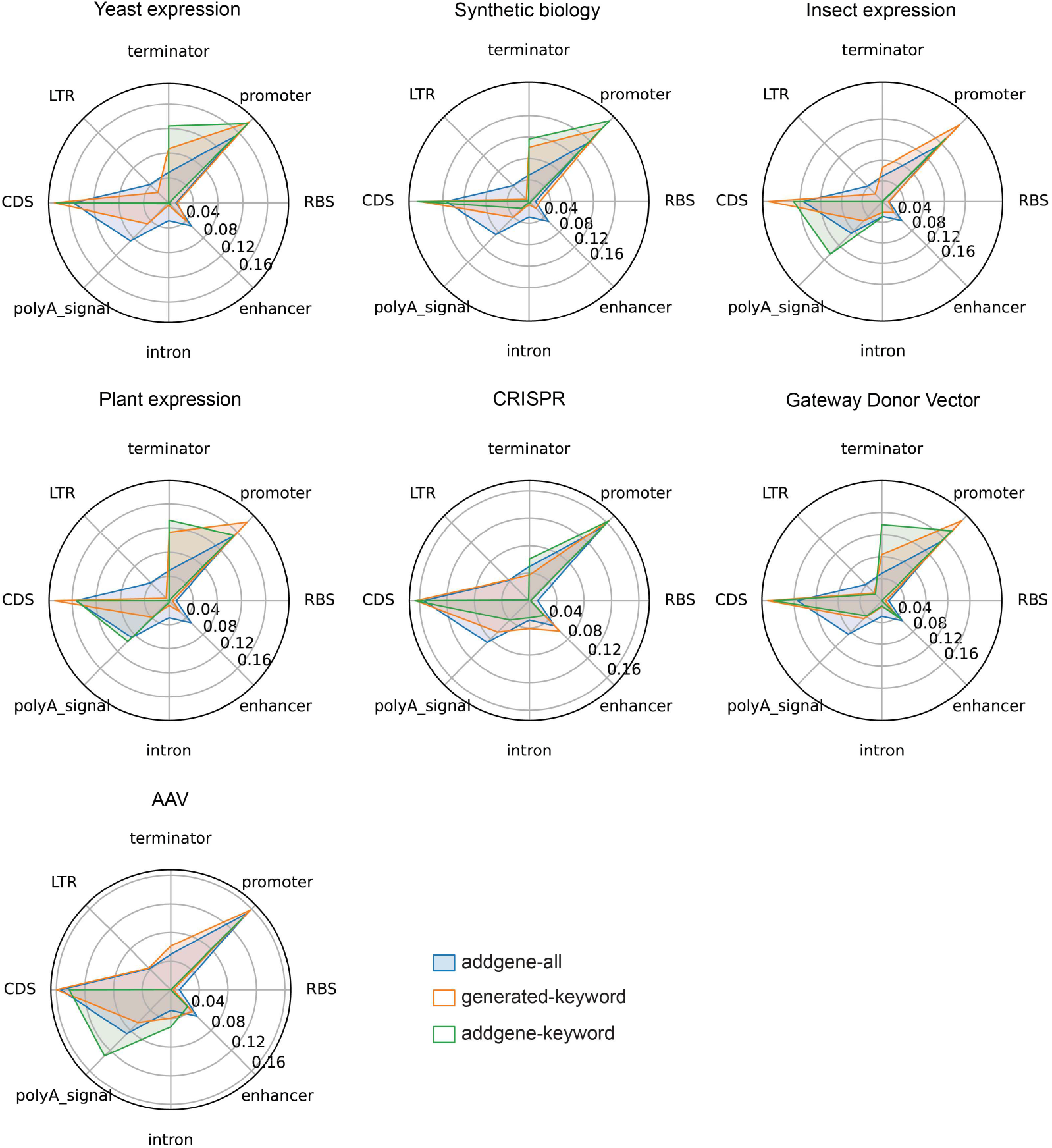
Part frequencies for plasmid sequences generated with the fine-tuned PlasmidGPT model. Part frequencies are shown for three subsets of sequences: randomly sampled training sequences (blue, n = 2,000), randomly sampled training sequences related to specific keywords (green, n = 200), and generated sequences with specific keywords (orange, n = 100).

**Supplementary Figure 4.**
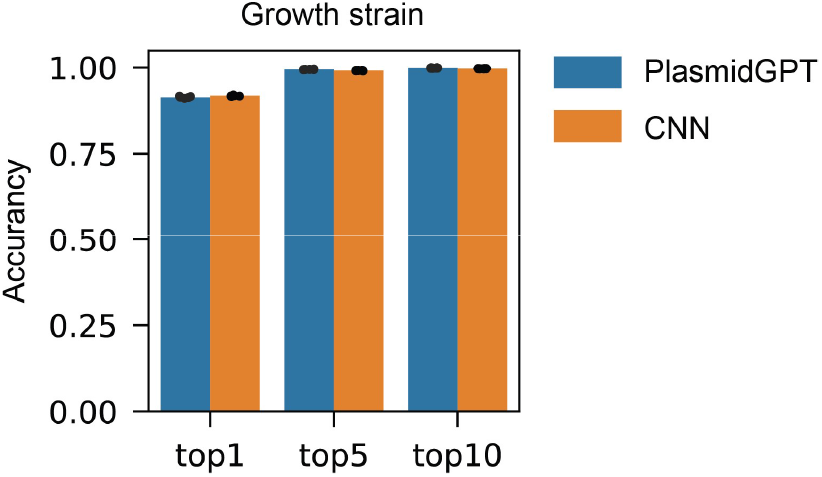
Performance of PlasmidGPT and CNN on the growth strain prediction task. We used 5-fold cross validation tests to evaluate model performance (n = 5) and error bars denote standard deviation.

**Supplementary Figure 5.**
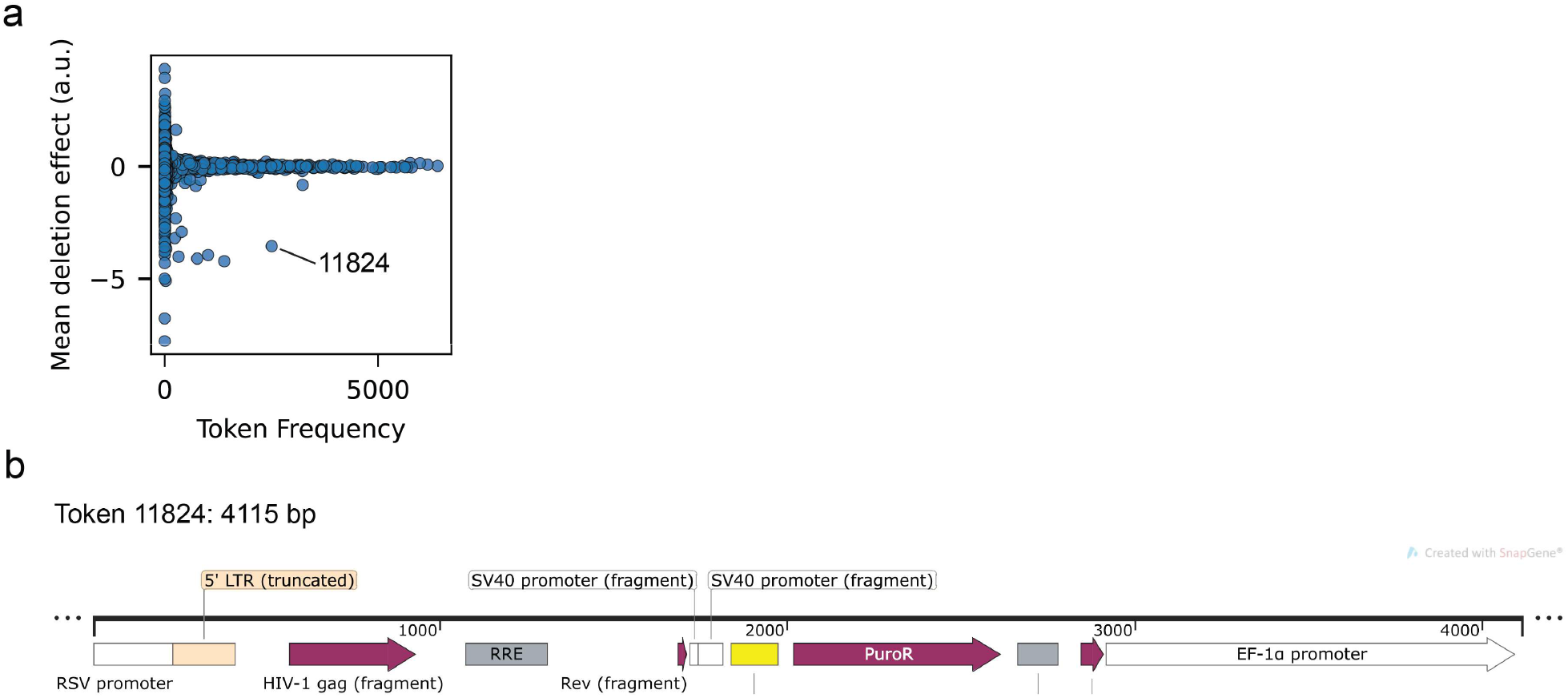
Deletion effect analysis of tokens for predicting Feng Zhang lab plasmids. **a**) Relationship of token frequency and mean deletion effect. Each dot represents the frequency and mean deletion effect for one token. The deletion effect was calculated by removing the token from the input sequence and comparing the activity of the output neuron with that of the unperturbed sequence. **b)** Annotation of token 11824 by pLannotate (4155 bp). Genetic parts are presented in different colors and the annotation map was generated using SnapGene Viewer.

**Supplementary Figure 6.**
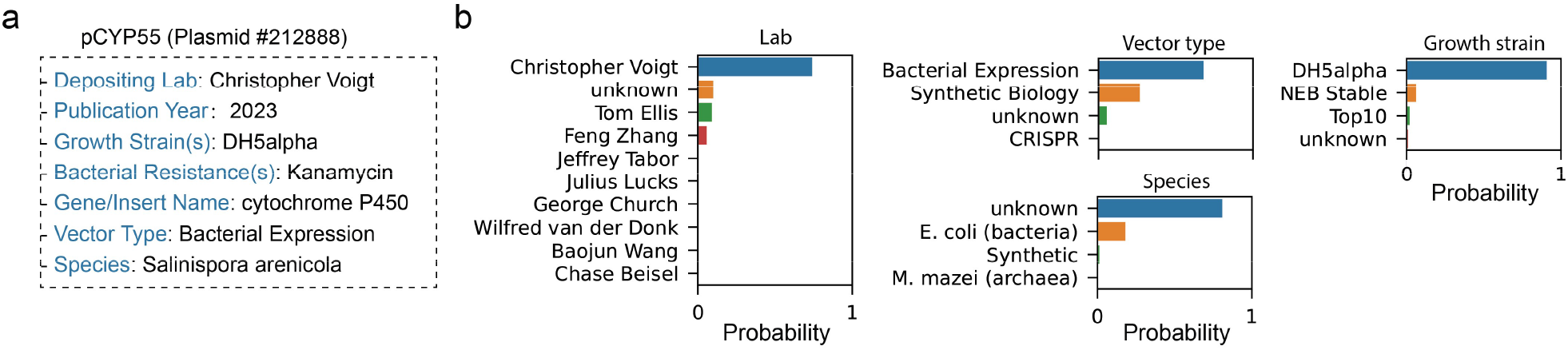
Annotation of the out of the distribution (OOD) plasmid. **a)** Plasmid #212888 was used as an example and the embeddings from the PlasmidGPT model were utilized to predict the attributes related to this plasmid. **(b)** Top predictions for each attribute.

**Supplementary Figure 7.**
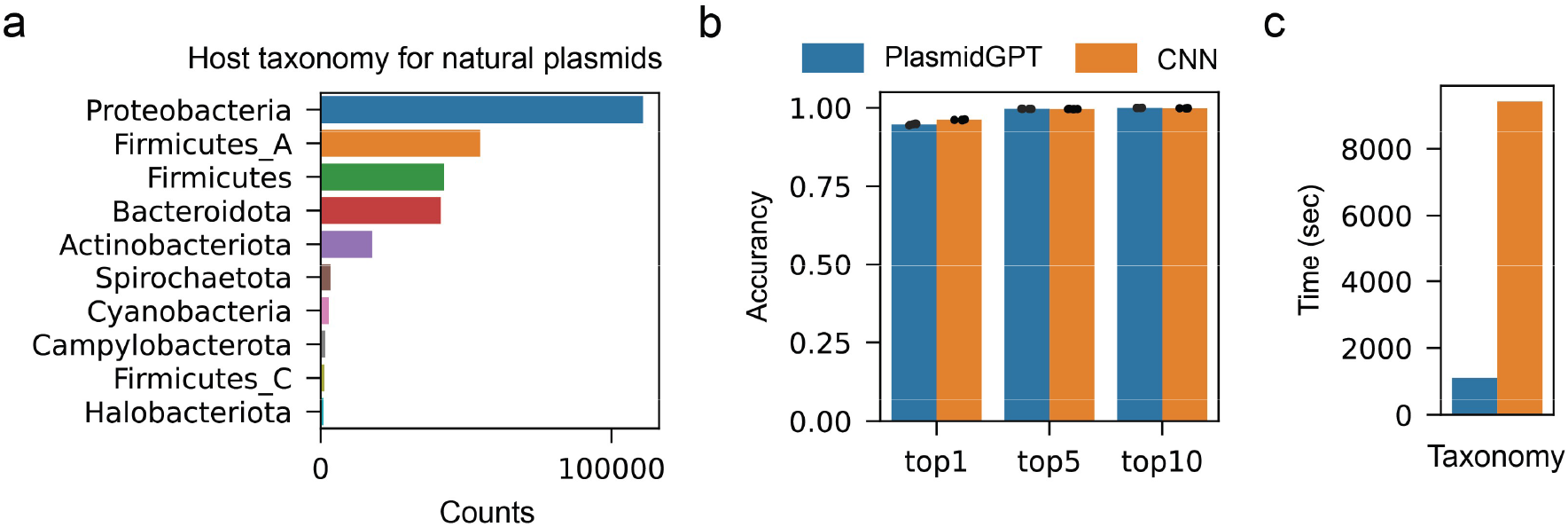
Prediction of host taxonomy for natural plasmids. **a)** Top 10 host taxonomies for the natural plasmids from the IMG/PR dataset^17^. **b)** Performance of PlasmidGPT and the CNN model on the host taxonomy prediction task. We used 5-fold cross validation tests to evaluate model performance (n = 5) and error bars denote standard deviation. **c)** Time consumption for the two models.

**Supplementary Table 1.**
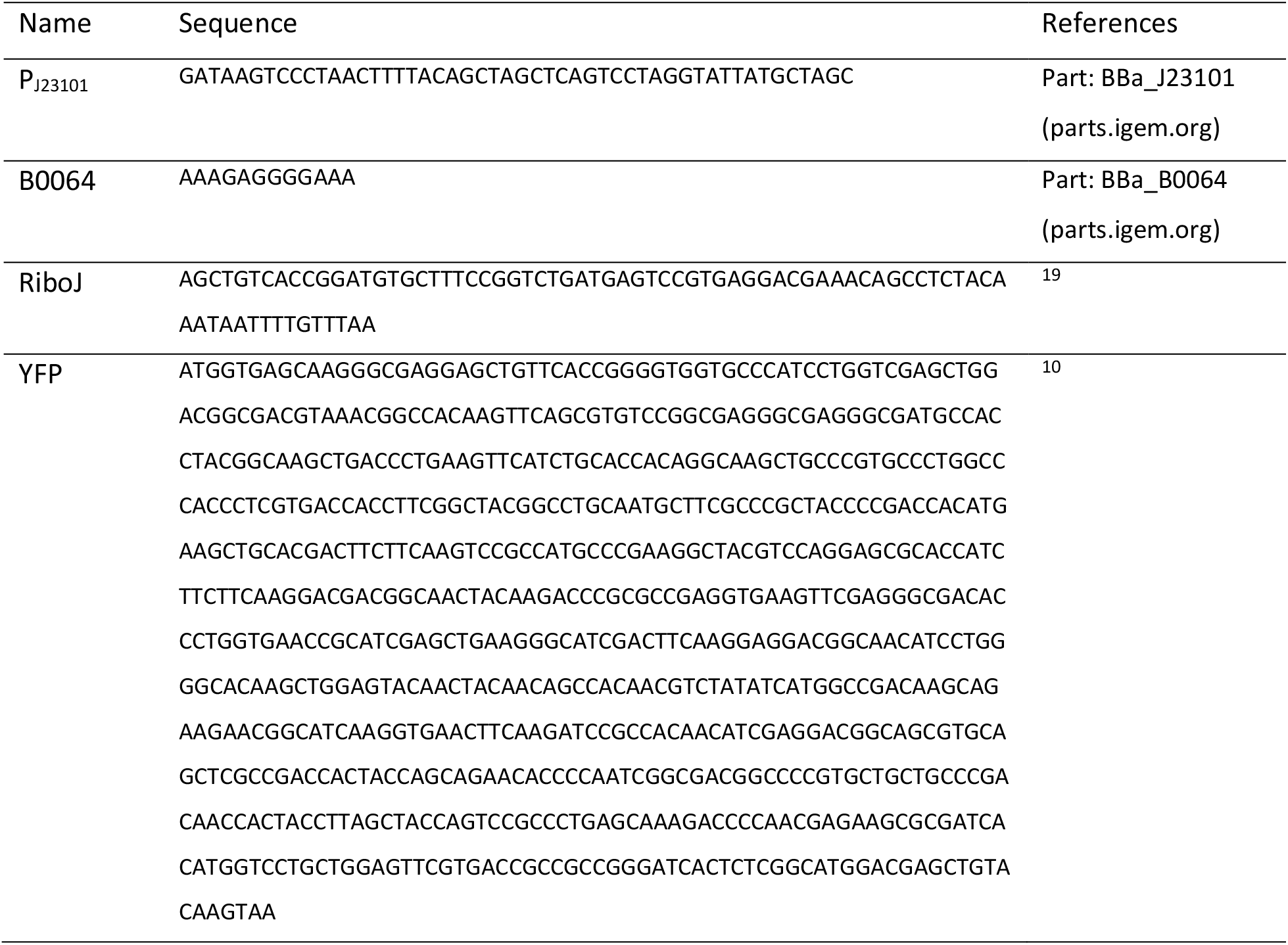
Sequences used in this study. The YFP expression cassette contains the following genetic parts.

